# Darwin’s small and medium ground finches might have taste preferences, but not for human foods

**DOI:** 10.1101/2021.07.17.452789

**Authors:** D. Lever, L. V. Rush, R. Thorogood, K. M. Gotanda

## Abstract

Urbanization is rapidly changing ecological niches. On the inhabited Galapagos Islands, Darwin’s finches consume human-introduced foods preferentially; however, it remains unclear why. Here we presented pastry with flavour profiles typical of human foods (oily, salty, sweet) to small ground finches (*Geospiza fuliginosa*) and medium ground finches (*Geospiza fortis*) to test if latent taste preferences might drive selection of human foods. If human-food flavours were consumed more than a neutral or bitter control only at sites with human foods, then we predicted tastes were acquired after urbanization; however, if no site-differences were found then this would indicate latent taste preferences. Contrary to both predictions, we found little evidence that human-food flavours were preferred compared to control flavours at any site. Instead, finches showed a weak aversion to oily foods, but only at remote (no human foods present) sites. This was further supported by behavioural responses, with beak wiping occurring more often at remote sites after finches tasted flavours associated with human foods. Our results suggest, therefore, that while Darwin’s finches may have acquired a tolerance to the flavours of human food, latent taste preferences are unlikely to have played a major role in their dietary response to increased urbanization.

## Introduction

Human behaviour is now recognised to be a strong driver of local adaptation and differences among populations of animals [1,2]. Urbanization [3–5], for example, can have profound effects on foraging because humans often introduce novel foods to the surrounding environment either intentionally (e.g. via garden bird feeding [6–8]) or unintentionally (e.g. by planting ornamental, invasive plants [9,10] or food waste [11,12]). This changes the diversity and availability of food items, and generates different foraging landscapes from those in which most animals evolved [13–15]. However, organisms’ responses to these altered niches vary, with some birds, for example, not only adapting more readily to incorporate human-foods into their diet, but even preferentially consuming them over native food sources [14,16–19]. These differences in whether, and how, populations and species utilise human foods can then have consequences for local adaptation [7,20] and potentially affect species divergence [21]. Taste preferences can also have implications in conservation, mediating the potential for invasive species to establish [22,23], for example, or making native species vulnerable to accidental poisoning [24]. Furthermore, the gut microbiome, which is increasingly recognised to affect a suite of physiological, immune, and cognitive functions in wild non-human animals [25,26], can also vary with consumption of human foods [27–29]. Nevertheless, understanding why some species in urban areas shift their diets to preferentially forage on human foods remains unclear.

Taste plays an important role in foraging as it allows individuals to detect nutritious versus unprofitable substances in food items [30,31]. Although taste was long assumed to be of little importance for birds, they have sophisticated sensory adaptations with taste receptors identified for bitter [30,32,33], sweet [30], and salt [34] flavours. Furthermore, experiments have demonstrated that birds can use bitter tastes to avoid toxic foods [32,34,35], use sweet tastes to detect sugars [30,34,36–39] indicating high caloric content [40], and use salty tastes to detect salts [39,41,42] indicative of necessary minerals and proteins [43]. Preferences for certain tastes can therefore evolve when flavours are linked to food quality and nutrient content [30]. For example, omnivorous and frugivorous birds are able to detect sugars at low levels and prefer sweet tastes compared to birds that forage on other types of food [34], potentially because this facilitates optimal foraging for caloric content of nectar and ripe fruit.

Taste preferences can also be maladaptive, however, if they lead to the preferential consumption of human foods. The introduction of human foods in urban areas causes changes in resource availability, including calories and nutrient concentration [44,45] as much of the human foods accessible to birds consists of discarded ‘junk food’ and snacks that are high in fat, sugar, and salt. The availability of these foods can alter natural nutritional landscapes with detrimental effects [24,46]. For example, hibernation in mammals is perturbed when human foods are consumed [47,48], and racoons feeding on human foods in urbanised areas show greater weight and blood sugar content [49]. In birds, American crow (*Corvus brachyrhynchos*) chicks are smaller [50] and Canadian geese (*Branta canadensis maxima*) have higher rates of angel wing disorder in urbanised areas, likely as a result of nutrient deficiency [51], and Australian magpies (*Gymnorhina tibicen*) show alterations to blood chemistry following backyard provisioning [52]. If animals are adapted to detect and prefer foods based on taste profiles that are coincidentally elevated in human foods, then they could assess these foods erroneously as high quality and favour consumption [24,45]. On the other hand, if their taste receptors are adapted to prefer flavours characteristic of human foods, then they might be better able to take advantage of this newly available resource if native foods decline (i.e. poor condition is preferable to starvation). Therefore, preferential consumption of different foods due to taste preferences, especially in the context of urbanization, could be either adaptive or maladaptive. However, it remains largely unknown whether taste preferences actually underlie the preferential consumption of human foods in urban areas.

Here, we investigate if Darwin’s finches, which on human-inhabited islands are known to vary in their preferential consumption of human foods [14], have taste preferences for flavours associated with human foods. Darwin’s finches are a model system for demonstrating how foraging ecology shapes adaptation into different ecological niches and results in an adaptive radiation [53,54]. Contemporary work has shown that the presence of humans is affecting traits such as beak morphology [55] and this has been linked to the consumption of human foods [21]. Indeed, we know that Darwin’s finches on human-inhabited islands can preferentially consume human foods including crisps (potato chips), biscuits (hard cookies), and rice at sites where these foods are abundant such as at tourist beaches and in urban areas [14]. Indeed, urban finches can have higher nesting success than non-urban finches [56], suggesting that at least some species of Darwin’s finches can become locally adapted to urban environments. Thus, the Darwin’s finches on the Galapagos represent an excellent opportunity to test whether taste preferences might underpin consumption of human foods where these are available and abundant, such as in urban or tourist areas.

We assessed taste preferences in Darwin’s finches in separate populations with little to no movement of finches among sites [57] that varied in the availability of human foods previously shown to be attractive to finches [14]. We presented flavours in the absence of visual cues typical to human foods to exclude the role of learned associations with packaging (e.g. crisp packets [14]). If taste preferences arise because of the consumption of human foods, we predicted finches at sites with human foods would only prefer flavours associated with human foods (e.g. sweet, salty, and oily) at sites where human foods are abundant. If taste preferences are innate (i.e. latent), we predicted preferences would occur for such flavours across sites, regardless of the presence or absence of human foods. Potential taste preferences can be quantified as variation in feeding rate, or as variation in post-feeding beak-wiping rate [30,58–60]. Although beak-wiping can clean the beak of debris [61], vigorous beak-wiping commonly occurs after feeding on something unpalatable [30,60]. Therefore, we measured taste preferences in terms of consumption and beak-wiping behaviour compared to two controls, neutral pastry that had no flavour added and bitter-flavoured pastry as a negative control. Bitter substances are aversive to many bird species, so we expected finches to consume less bitter-flavoured pastry compared to neutral or other flavoured pastry across all sites as well as to elicit more beak-wiping behaviour [30,60].

## Materials and Methods

### Study species and location

We focused on Darwin’s finches, an endemic group of passerines on the Galapagos Islands, at three sites on Santa Cruz Island that varied in their exposure to human foods (Supplemental Figure 1). The remote site was a non-urban site 12 km from the main urban town with no presence of human foods (El Garrapatero proper) [27], the beach site was El Garrapatero beach, a tourist, non-urban site 12 km from Puerta Ayora where visitors often bring picnics so human food is present and abundant [14,27], and the town site was Puerto Ayora, a fully urbanized town where humans and their food are ubiquitous throughout the entire town [14]. The two focal species were small ground finches (*Geospiza fuliginosa*) and medium ground finches (*Geospiza fortis*). Galápagos mockingbirds (*Mimus parvulus*) and two other finch species (the cactus finch, *Geospiza scandens* and the small tree finch, *Camarhynchus parvulus*) were also present occasionally, but rarely interacted with our experiments.

### Experimental protocol and video recording

We conducted trials at each of the three sites (remote = 16 trials, beach = 15 trials, and town = 18 trials; Supplemental Table 1) using a ‘cafeteria’ tray experiment [14]. Five plastic cups with the diameter of a large chicken egg were randomly positioned in the periphery of a 3×3 egg carton and placed on the ground on a white plastic dinner plate that was visible below the egg carton (e.g. the carton did not hang over the sides of the plate; Supplemental Figure 2). The central dimple of the egg carton was weighted with a small rock, and the unused dimples were left empty. The trial began when the first bird approached the tray and fed, and then continued for 10 minutes [14]. If no finches fed, the trial was aborted after 20 minutes. All trials were performed during the rainy season from February 20^th^ to March 25^th^, 2018, between 6am and 11am, or 3pm and 6pm and were filmed using a video camera (Sony HDR-CX625 Full HD Compact Camcorder or Canon 7D Mark II with 100-400mm lens) positioned 10 metres from the cafeteria tray. The majority of individuals were not uniquely identifiable (fewer than 4%), so we cannot be sure that birds participating in different trials were independent. However, to reduce the potential for pseudo-replication between trials, we conducted each trial at least 100 m apart within the study locations.

Each cup was filled with 2.5 g of pastry made from flour, unsalted butter, and water, following methods from Speed and colleagues (2000). The pastry (335 g flour, 135 g unsalted butter, and 30 g water) was flavoured according to commonly available human-foods in the environment [14] and each cup was coloured (blue, green, pink, purple, and yellow) to facilitate recognition of the contents: (i) blue indicated high in fat (6g vegetable oil/pastry batch), (ii) green indicated bitter (0.1g quinine/pastry batch), (iii) purple indicated sweet (23g sugar/pastry batch), (iv) yellow indicated salty (1.333g salt/pastry batch), and (v) pink indicated neutral or unflavoured pastry. To habituate the birds to the experimental set-up, we first conducted trials at each site with only unflavoured pastry (remote = 17 trials, beach = 17 trials, town = 19 trials; Supplemental Materials, Supplemental Table 1). Birds can have latent colour preferences, either from experience or evolutionary history [e.g., 62]. However, we detected no strong biases within finch species towards, or against, any of the coloured cups based on these trials (Supplemental Materials; Supplemental Table 2; Supplemental Figures 3 & 4). Therefore, any preferences detected using flavoured pastry were most likely due to taste and not visual preferences.

### Video analysis

Videos were analysed using BORIS (Behavioral Observation Research Interactive Software) (Friard and Gamba, 2016) and each 10-minute trial was analysed by one observer (DL). Species identification was done based on bill and body size in comparison to the size of the cafeteria tray. The observer was trained by KMG to first identify still images of birds (taken from videos collected during trials), and then by using the videos. We counted the number of feeding events at the level of the trial and assigned these to each species of finch. We defined a feeding event as when a bird’s beak was submerged into a cup, lifted, and then food was consumed (Supplemental Figure 2). Following each feeding event, we then recorded the number of times the finch wiped its beak on a surface within 20 seconds in accordance with published methods on beak-wiping [60,61].

### Statistical analyses

Statistical analyses were undertaken using the R environment version 4.0.2 (R Core team, 2020; data and code for analyses are available as Supplemental Materials). To analyse differences in taste preferences, we used generalised linear mixed-effect models (GLMMs) with a negative binomial error distribution (using the glmer.nb() function in the lme4 package; [63] to account for overdispersion in the number of feeding events per trial (response variable). Species nested within trial was included as a random effect to account for non-independence of feeding events, and the fixed effects were species, site, and pastry flavour. The reference level (i.e. the model intercept) was ‘neutral flavour (pink)’ and ‘town’. We included an interaction between site and pastry flavour, and assessed whether it contributed significantly to model fit using a likelihood ratio test (compared to a simpler model with the same random effect structure, but containing only additive fixed effects). We then used z-tests to assess the significance of differences in consumption among pastry flavours. We report estimates and standard errors and provide incidence rate or odds ratios (negative binomial and binomial models, respectively) to compare effects.

The number of beak wipes following a feeding event were low (5 or fewer) and highly right-skewed (medium ground finch = 3.69, small ground finch = 3.44; calculated using the ‘moments’ package; [64] so we therefore modelled the occurrences of beak wipes using a binomial distribution, where the denominator in the response variable was the number of feeding events when no beak wipes occurred. Assumptions of homogeneity of variance and uniformity of the residuals for all models were checked using Kolmogorov-Smirnov tests for uniformity, simulation tests for dispersion, and a binomial test for outliers (implemented using the ‘DHARMa’ package [65]). The cactus finch and small tree finch rarely came to the experimental trays so only medium and small ground finches were included in the dataset (see Supplemental Figures 3 & 4).

## Results

A total of 49 taste preference trials were conducted across three sites. Participation was similar for both species (medium ground finch = 16 remote trials and 10 town trials, small ground finch = 10 remote trials and 14 town trials) except for trials conducted at beach sites where medium ground finches participated in only 4 of the 15 trials whereas small ground finches participated in all trials (Supplemental Table 1).

### Taste preferences

The overall consumption of pastry among ground finch species did not differ (1089 feeding events by medium ground finches versus 1186 feeding events by small ground finches; estimate = −0.102 ± 0.163, z = −0.625, p = 0.532). However, we found some evidence that ground finches preferred some flavour types more than others at different sites (flavour-type * site, χ^2^ = 16.352, d.f. = 8, p = 0.0376; Figure 1A & 1B; Supplemental Table 3). To investigate this interaction further, we explored differences among flavour-types in separate models for each site. At beach sites, there was no significant difference in the number of feeding events according to flavour-type (χ^2^ = 2.025, d.f. = 4, p = 0.731, N = 15 trials). However, at remote (χ^2^ = 20.090, d.f. = 4, p = 0.0005, N = 16 trials) and town (χ^2^ = 10.595, d.f. = 4, p = 0.032, N = 18 trials) sites we detected significant variation in consumption and the preferred flavours differed. Ground finches at remote sites fed less often on oily (blue) pastry than either the neutral or the bitter control whereas at sites in town, finches fed more often on sweet (purple) pastry, but only in comparison to the bitter control (Table 1, Figure 1A & 1B).

**Table 1.** Mean differences (± S.E.) in the (I) number of feeding events and (II) proportion of feeding events followed by beak-wiping at (a) Town sites, (b) Beach sites, and (c) Remote sites that vary in exposure to human foods. Medium ground finches (I: N = 130 observations from 26 trials, II: N = 95 observations from 26 trials) and small ground finches (I: N = 195 observations from 39 trials, II: N = 114 observations from 39 trials) were presented with coloured cups containing pastry flavoured to be oily (blue), sweet (purple), or salty (yellow), and two controls: neutral (pink, set as model intercept) and bitter (green). Differences in (I) were estimated using generalised linear mixed effects models with a negative binomial error distribution and where trial and site were included as random effects. Differences in (II) were estimated using a similar model but with a binomial error distribution to account for proportional response data. Significant differences are indicated in bold font.

|  | I. Feeding<br>events |  |  | II. Beak-wiping |  |  |
| --- | --- | --- | --- | --- | --- | --- |
|  | Mean difference<br>± S.E. | Z | p | Mean<br>difference ±<br>S.E. | Z | p |
| (a) Town sites: |  |  |  |  |  |  |
| intercept(neutral) | 1.872 ± 0.290 | 6.447 | <0.001 | -2.696 ± 0.345 | -7.489 | <0.001 |
| Oily (blue) | 0.118 ± 0.347 | 0.340 | 0.734 | 0.905 ± 0.433 | 2.090 | 0.146 |
| Bitter (green) | -0.435 ± 0.345 | -1.262 | 0.207 | -0.053 ± 0.403 | -0.131 | 0.393 |
| Sweet (purple) | 0.519 ± 0.397 | 1.310 | 0.190 | 0.936 ± 0.368 | 2.541 | 0.070 |
| Salty (yellow) | -0.591 ± 0.348 | -1.699 | 0.089 | 0.593 ± 0.399 | -1.486 | 0.369 |
| Species(small) | 0.446 ± 0.236 | 1.888 | 0.059 | 0.565 ± 0.320 | -0.180 | 0.857 |
| (b) Beach sites: |  |  |  |  |  |  |
| intercept(neutral) | 3.120 ± 0.554 | 5.635 | <0.001 | -1.396 ± 0.429 | -3.255 | 0.001 |
| Oily (blue) | -0.300 ± 0.513 | -0.582 | 0.560 | **-0.820 ± 0.414** | **-1.980** | **0.048** |
| Bitter (green) | 0.018 ± 0.477 | 0.038 | 0.970 | **-0.733 ± 0.347** | **-2.112** | **0.035** |
| Sweet (purple) | -0.249 ± 0.492 | -0.506 | 0.613 | -0.647 ± 0.375 | -1.725 | 0.085 |
| Salty (yellow) | -0.512 ± 0.463 | -1.107 | 0.268 | -0.030 ± 0.341 | -0.089 | 0.929 |
| Species(small) | -0.348 ± 0.418 | -0.832 | 0.406 | 0.253 ± 0.352 | 0.720 | 0.471 |
| (c) Remote sites: |  |  |  |  |  |  |
| intercept(neutral) | 2.241 ± 0.240 | 9.328 | <0.001 | -2.969 ± 0.324 | -9.172 | <0.001 |
| Oily (blue) | **-0.766 ± 0.291** | **-2.638** | **0.008** | **0.874 ± 0.436** | **2.006** | **0.045** |
| Bitter (green) | 0.328 ± 0.270 | 1.216 | 0.224 | 0.210 ± 0.380 | 0.553 | 0.580 |
| Sweet (purple) | 0.427 ± 0.271 | 1.575 | 0.115 | **1.314 ± 0.334** | **3.915** | **<0.001** |
| Salty (yellow) | -0.033 ± 0.276 | -0.119 | 0.905 | **0.982 ± 0.365** | **2.689** | **0.007** |
| Species(small) | -0.385 ± 0.247 | -1.558 | 0.119 | **1.020 ± 0.240** | **4.257** | **<0.001** |

**Figure 1.**
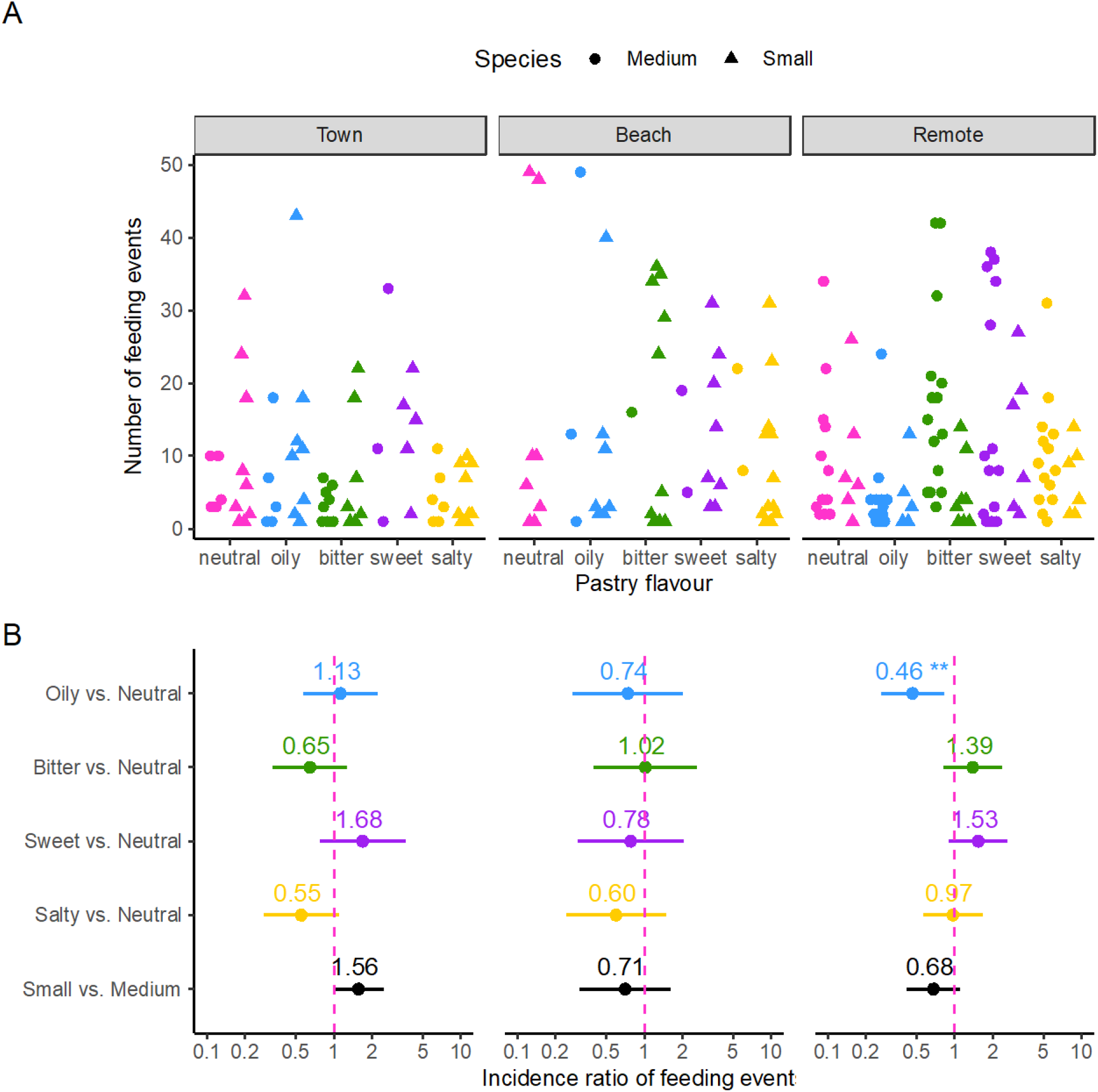
Differences in the number of feeding events by medium and small ground finches presented with coloured cups containing pastry with neutral (pink), oily (blue), bitter (green), sweet (purple), or salty (yellow) flavours at either town (N = 18 trials), beach (N = 15 trials), or remote (N = 16 trials) sites. (A) presents the raw data, (B) presents the effect sizes of the differences between each flavour and the neutral pastry, or between species, computed from generalised linear mixed effects models (see Methods for more details). Effects significantly different from zero (dashed pink vertical line) are indicated by asterisks (* 0.01 < p < 0.05, ** 0.001 < p < 0.01).

We next assessed behavioural wiping responses to each flavour-type (Figure 2). Overall, small ground finches were much more likely to wipe their beak after feeding than medium ground finches (estimate = 0.528 ± 0.198, z = 2.672, p = 0.008; Supplemental Table 4). Again, however, we detected that beak-wiping was more likely to occur after consuming some flavours at some sites (flavour-type * site, χ^2^ = 28.132, d.f. = 8, p = 0.0005; Figure 2; Supplemental Table 4). On further inspection (as above), at town sites ground finches showed no difference in their propensity to wipe their beak after feeding on flavoured pastry (Table 1, Figure 2, N = 18 trials). However, at beach sites ground finches wiped their beaks less often after consuming oily (blue) or bitter (green) flavoured pastry compared to neutral (pink) pastry and less often after eating salty (yellow) compared to bitter pastry (Table 1, Figure 2, N = 15 trials). At remote sites, ground finches were more likely to wipe their beaks after consuming pastry containing any ‘human-food’ flavour as compared to either the neutral or the bitter control (Table 1, Figure 2, N = 16 trials). Taken together, these results suggest that ground finches in town showed little preference to human-food flavours whereas at remote sites, all human-food flavours evoked more behavioural reactions than controls, and oily flavoured pastry was preferred the least.

**Figure 2.**
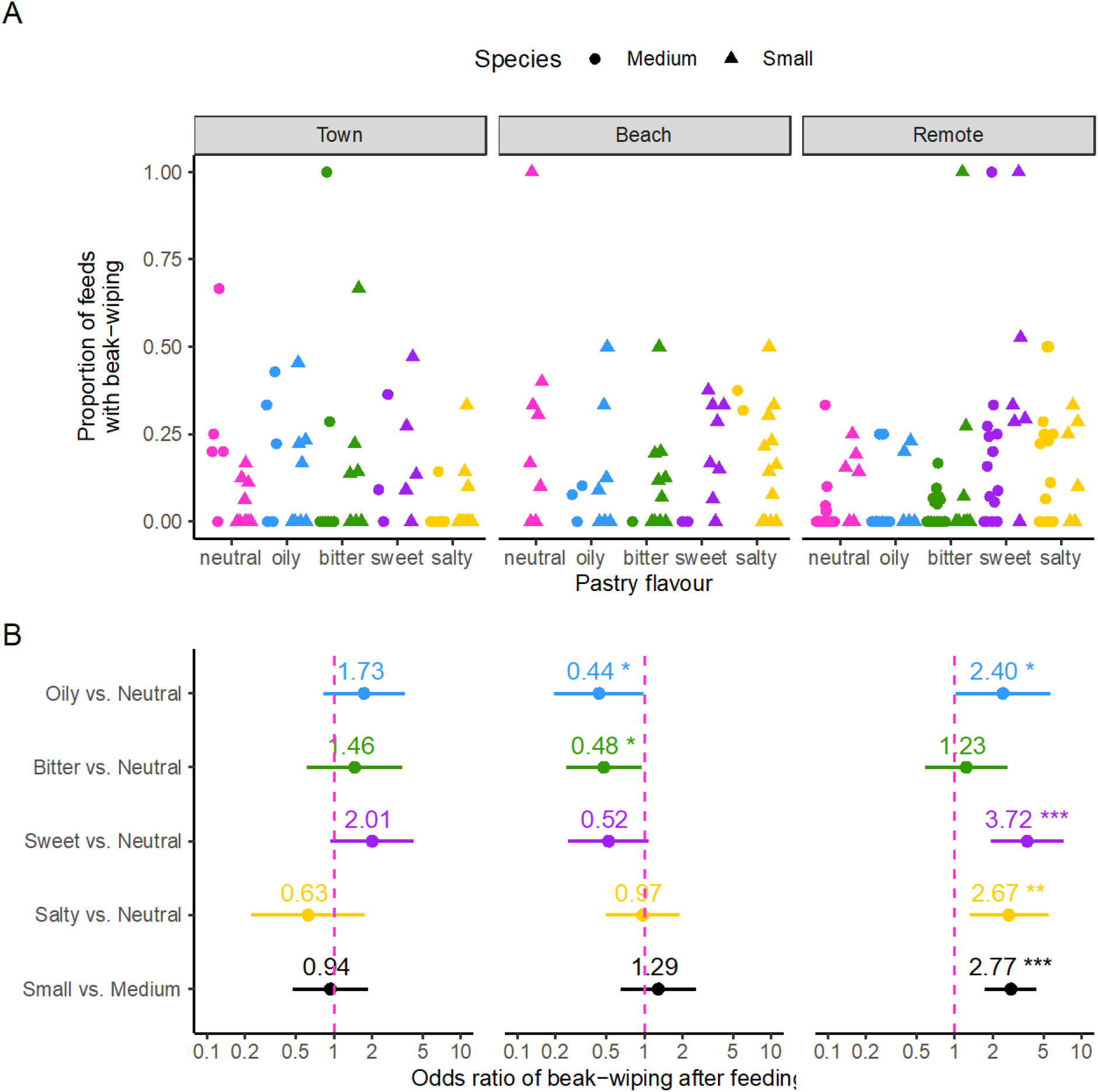
Differences in the proportion of feeding events that were followed by beak-wiping by medium and small ground finches presented with coloured cups containing pastry with neutral (pink), oily (blue), bitter (green), sweet (purple), or salty (yellow) flavours at either town (N = 18 trials), beach (N = 15 trials), or remote (N = 16 trials) sites. (A) presents the raw data, (B) presents the effect sizes of the differences between each flavour and the neutral pastry, or between species, computed from generalised linear mixed effects models (see Methods for more details). Effects significantly different from zero (dashed pink vertical line) are indicated by asterisks (* 0.01 < p < 0.05, ** 0.001 < p < 0.01).

## Discussion

Taste preferences have often been overlooked in understanding animals’ foraging decisions, yet in human-modified environments, latent taste preferences could explain why some species are able to readily adapt to novel foods while others do not. Here we investigated if taste preferences can explain preferential consumption of human foods by Darwin’s finches [14]. We predicted that if finches at sites with more exposure to human foods (i.e. at the tourist beach or in town) showed greater consumption and reduced aversive behavioural responses to flavours typical of these foods (salty, oily, sweet), then taste preferences could have developed from experience with the changed foraging landscape. However, if finches across sites preferred these ‘human-food flavours’ then latent taste preferences could have facilitated rapid adoption of human foods into the diet. Against both predictions, however, the only evidence we detected for a taste preference was that ground finches at remote sites showed a weak aversion to oily foods. This was further supported by behavioural responses, with beak wiping (an aversive response common across birds [30,60]) occurring more often at remote sites after finches tasted flavours associated with human foods. This suggests that ground finches do not have latent taste preferences for human foods or have acquired preferences from contact with human foods. However, ground finches might now be more tolerant of oily flavours. It is possible that our sample sizes were too small to detect preferences for human-food flavours, or that we did not add sufficient flavour to the pastry for these to be detectable. However, previous work detecting taste preferences had smaller sample sizes than our study (e.g., n = 6 per group [36]; n = 11 [37]; n = 6 and 10 [38]), and the amount of constituents that we added to the pastry emulated human foods as closely as possible. It therefore seems unlikely that our results can only be explained by methodological issues. Since Darwin’s finches consume human foods preferentially when available, why did we not find preferences for flavours associated with commonly available human foods?

It could be that Darwin’s finches have not evolved a preference for tastes associated with human foods because these species are generalist feeders, especially during periods of non-drought [13]. We conducted our trials during the traditional rainy season when the finches would be more generalist. Taste preferences evolve when they allow animals to identify food items that offer important nutrients (e.g., high in lipids, salts, or sugars)[30,31,34,46]. Yet for generalists, it might not be adaptive to have latent taste preferences if these limit individuals from consuming a wide variety of dietary items [13] or they might not need to discern specific foods that are high in lipids, salts, or sugars. Indeed, many studies that have found taste preferences in birds have been conducted with specialists [37,38]. Another possibility is that Darwin’s finches have not yet acquired preferences for flavours associated with human foods. Tourism has grown exponentially only relatively recently [66]. At the tourist beach site, easy access to the public only became available around 2010 [J. Podos, personal communication], and the town of Puerto Ayora was established in 1926 [67; E. Hennessey, personal communication], so perhaps not enough generations have passed from when finches gained access to human foods for finches to acquire taste preferences.

For the finches, we found that the presence or absence of human foods did not correlate with different taste preferences as predicted. Surprisingly, we also found little evidence that the bitter-tasting control elicited beak wiping. This was unexpected because we know birds possess TAS2R bitter taste receptors [33], and often discriminate against toxic prey via bitterness. In fact, of the species studied by Wang and Zhao [33], medium ground finches had the second most TAS2R genes. So why did we not find a lack of aversion to bitter tastes? If the natural foods found at remote sites are bitter in taste, then perhaps finches are accustomed to bitter tastes. Bitter tastes are often associated with aposematic prey, and the relative lack of aposematic prey on the Galapagos [68] suggests finches may not need to discriminate against bitter tastes. Another possibility is zoopharmacognosy, where animals eat medicinally advantageous foods, despite possible aversive qualities. This is common in birds; great bustards (*Otis tarda*) ingest toxic blister beetles to control digestive tract parasites [69], and house sparrows (*Passer domesticus*) ingest leaves containing quinine (our bittering agent) during malaria outbreaks [70], alleviating symptoms. Quinine is an invasive plant found on the Galapagos. However, no finch has ever been observed consuming quinine (Heinke Jäger, personal communication), so this possibility is unlikely.

The Galapagos Islands are experiencing an exponential increase in urbanization and tourism, including permanent human residents [66], and we know Darwin’s finches preferentially consume human foods over natural food sources when readily available [14]. However, here we found little evidence for taste preferences for flavours associated with human foods during the traditional rainy season, although finches at remote sites showed a small aversion towards oily foods. It therefore seems likely that finches do not have latent preferences for these flavours, nor acquired a preference through repeated exposure to human foods. Why then have Darwin’s finches adapted rapidly to changing food availability and incorporated human foods into their diet when human foods are readily available? One possibility is that they could be attracted to other sensory cues such as aural or visual cues associated with human foods. For example, in town and on the beach (but not in remote areas), finches respond to brightly coloured visual cues of human food packaging and are attracted to the ‘crinkle’ sound associated with foil and plastic food packaging [Supplemental Figure 5,14]. Alternatively, it could be driven by availability itself at the beach and town sites. While the food sources Darwin’s finches normally feed upon are available within town and at the beach [14] (though we did not control for this), the abundance of human foods at these sites simply make these types of food more accessible to finches, and therefore, finches did not need to discriminate between different flavours typically associated with human foods and aversive flavours to expand their diet diversity [14]. Further work is required to understand the mechanisms underlying how Darwin’s finches developed a preference for consuming human foods at sites where human foods are readily available.

Humans, through processes such as urbanization, can have a major impact on foraging ecology by introducing novel foods that can become preferentially consumed by birds [16,17]. However, the mechanisms leading to changes in foraging ecology remain largely unknown. Although we cannot yet explain *why* Darwin’s finches prefer human foods, our results help to rule out the possibility that taste preferences play an important role in incorporating human foods into their diets. Similarly, our finding that ground finches do not find bitter tastes aversive expands the increasing knowledge on variation in response to tastes among species. As the adoption of human foods into animals’ diets can have cascading effects on health, reproduction, and fitness [6,15,20,47,49], it remains of paramount importance to elucidate why some species integrate these foods while others do not.

## Author contributions

KMG and RT designed the study, KMG and LVR collected the field data, DL analyzed the videos, RT, DL, and KMG analyzed the data, and all authors wrote and edited the manuscript.

## Acknowledgements

We thank Lotte Skovmand for their assistance with field work and Nick Davies for their assistance with the research project. During this study, RT was supported by an Independent Research Fellowship from the Natural Environment Research Council UK (NE/K00929X/1) and a start-up grant from the Helsinki Institute of Life Science (HiLIFE), University of Helsinki. KMG was funded by Christ’s College and Clare Hall at the University of Cambridge and was supported by Banting Postdoctoral Fellowship from the Natural Sciences and Engineering Research Council of Canada.

## References

1. Hendry AP, Gotanda KM, Svensson EI. 2017 Human influences on evolution, and the ecological and societal consequences. Philosophical Transactions of the Royal Society B 372, 20160028. (doi:10.1098/rstb.2016.0028)

2. Sullivan AP, Bird DW, Perry GH. 2017 Human behaviour as a long-term ecological driver of non-human evolution. Nature Ecology & Evolution 1, 0065. (doi:10.1038/s41559-016-0065)

3. Johnson MTJ, Munshi-South J. 2017 Evolution of life in urban environments. Science 358, eaam8237. (doi:10.1126/science.aam8327)

4. Alberti M, Marzluff J, Hunt VM. 2017 Urban driven phenotypic changes: empirical observations and theoretical implications for eco-evolutionary feedback. Philosophical Transactions of the Royal Society B 372, 20160029. (doi:10.1098/rstb.2016.0029)

5. Alberti M, Correa C, Marzluff JM, Hendry AP, Palkovacs EP, Gotanda KM, Hunt VM, Apgar TM, Zhou Y. 2017 Global urban signatures of phenotypic change in animal and plant populations. Proceedings of the National Academy of Sciences 114, 8651–8956. (doi:10.1073/pnas.1606034114)

6. Shutt JD, Trivedi UH, Nicholls JA. 2021 Faecal metabarcoding reveals pervasive long-distance impacts of garden bird feeding. Proceedings of the Royal Society of London B-Biological Sciences 288, 20210480. (doi:10.1098/rspb.2021.0480)

7. Badyaev A V, Young RL, Oh KP, Addison C. 2008 Evolution on a local scale: developmental, functional, and genetic bases of divergence in bill form and associated changes in song structure between adjacent habitats. Evolution 62, 1951–1964. (doi:10.1111/j.1558-5646.2008.00428.x)

8. Tryjanowski P et al. 2015 Urban and rural habitats differ in number and type of bird feeders and in bird species consuming supplementary food. Environmental Science and Pollution Research 22, 15097–15103. (doi:10.1007/s11356-015-4723-0)

9. Marco A, Lavergne S, Dutoit T, Bertaudiere-Montes V. 2010 From the backyard to the backcountry: how ecological and biological traits explain the escape of garden plants into Mediterranean old fields. Biological Invasions 12, 761–779. (doi:10.1007/s10530-009-9479-3)

10. Gelmi-Candusso TA, Hämäläinen AM. 2019 Seeds and the city: the interdependence of zoochory and ecosystem dynamics in urban environments. Frontiers in Ecology and Evolution 7, 1–19. (doi:10.3389/fevo.2019.00041)

11. Gajdon GK, Fijn N, Huber L. 2006 Limited spread of innovation in a wild parrot, the kea (*Nestor notabilis*). Animal Cognition 9, 173–181. (doi:10.1007/s10071-006-0018-7)

12. Auman HJ, Meathrel CE, Richardson A. 2008 Supersize me: does anthropogenic food change the body condition of Silver Gulls? A comparison between urbanized and remote, non-urbanized areas. Waterbords 31, 122–126.

13. De León LF, Podos J, Gardezi T, Herrel A, Hendry AP. 2014 Darwin’s finches and their diet niches: the sympatric coexistence of imperfect generalists. Journal of Evolutionary Biology 27, 1093–1104. (doi:10.1111/jeb.12383)

14. De León LF, Sharpe DMT, Gotanda KM, Raeymaekers JAM, Chaves JA, Hendry AP, Podos J. 2018 Urbanization erodes niche segregation in Darwin’s finches. Evolutionary Applications 12, 132–1343. (doi:10.1111/eva.12721)

15. Fuirst M, Veit RR, Hahn M, Dheilly N, Thorne LH. 2018 Effects of urbanization on the foraging ecology and microbiota of the generalist seabird *Larus argentatus*. PLoS One 13, e0209200. (doi:10.1371/journal.pone.0209200)

16. Sol D, Santos DM, Garcia J, Cuadrado M. 1998 Competition for food in urban pigeons: the cost of being juvenile. The Condor 100, 298–304. (doi:10.2307/1370270)

17. Marzluff JM, Neatherlin E. 2006 Corvid response to human settlements and campgrounds: causes, consequences, and challenges for conservation. Biological Conservation 130, 301–314. (doi:10.1016/j.biocon.2005.12.026)

18. Lowry H, Lill A, Wong BBM. 2013 Behavioural responses of wildlife to urban environments. Biological Reviews 88, 537–549. (doi:10.1111/brv.12012)

19. Tryjanowski P, Møller AP, Morelli F, Indykiewicz P, Zduniak P, Myczko Ł. 2018 Food preferences by birds using bird-feeders in winter: a large-scale experiment. Avian Research 9, 1–6. (doi:10.1186/s40657-018-0111-z)

20. Bosse M et al. 2017 Recent natural selection causes adaptive evolution of an avian polygenic trait. Science 358, 365–368. (doi:10.1126/science.aal3298)

21. De León LF, Raeymaekers JAM, Bermingham E, Podos J, Herrel A, Hendry AP. 2011 Exploring possible human influences on the evolution of Darwin’s finches. Evolution 65, 2258–2272. (doi:10.1111/j.1558-5646.2011.01297.x)

22. Snyder WE, Evans EW. 2006 Ecological effects of invasive arthropod generalist predators. Annual Review of Ecology, Evolution, and Systematics 37, 95–122. (doi:10.1146/annurev.ecolsys.37.091305.110107)

23. Shik JZ, Dussutour A. 2020 Nutritional dimensions of invasive success. Trends in Ecology & Evolution 35, 691–703. (doi:10.1016/j.tree.2020.03.009)

24. 2020 Kea cause of death confirmed. Department of Conservation Report

25. Davidson GL, Cooke AC, Johnson CN, Quinn JL. 2018 The gut microbiome as a driver of individual variation in cognition and functional behaviour. Philosophical Transactions of the Royal Society B 353, 20170286. (doi:10.1098/rstb.2017.0286)

26. Davidson GL, Raulo A, Knowles SCL. 2020 Identifying microbiome-mediated behaviour in wild vertebrates. Trends in Ecology & Evolution 35, 972–980. (doi:10.1016/j.tree.2020.06.014)

27. Knutie SA, Chaves JA, Gotanda KM. 2019 Human activity can influence the gut microbiota of Darwin’s finches in the Galapagos Islands. Molecular Ecology 28, 2441–2450. (doi:10.1111/mec.15088)

28. Littleford-Colquhoun BL, Frere CH, Weyrich LS, Kent N. 2019 City life alters the gut microbiome and stable isotope profiling of the eastern water dragon (*Intellagama lesueurii*). Molecular Ecologyecular Ecology 28, 4592–4607. (doi:10.1111/mec.15240)

29. Teyssier A, Matthysen E, Hudin NS, Neve L De, White J, Lens L. 2020 Diet contributes to urban-induced alterations in gut microbiota: experimental evidence from a wild passerine. Proceedings of the Royal Society B-Biological Sciences 287, 20192182. (doi:10.1098/rspb.2019.2182)

30. Rowland HM, Parker MR, Jiang P, Reed DR, Beauchamp GK. 2014 Comparative Taste Biology with Special Focus on Birds and Reptiles. In Handbook of Olfaction and Gustation, 4th Edition (ed RL Doty), pp. 957–982. Jon Wiley and Sons.

31. Skelhorn J, Halpin CG, Rowe C. 2016 Learning about aposematic prey. Behavioral Ecology 27, 955–964. (doi:10.1093/beheco/arw009)

32. Li D, Zhang J. 2013 Diet shapes the evolution of the vertebrate bitter taste receptor gene repertoire. Molecular Biology and Evolution 31, 303–309. (doi:10.1093/molbev/mst219)

33. Wang K, Zhao H. 2015 Birds generally carry a small repertoire of bitter taste receptor genes. Genome Biology and Evolution 7, 2705–15. (doi:10.1093/gbe/evv180)

34. Clark L, Hagelin J, Werner S. 2014 The Chemical Senses in Birds. In Sturkie’s Avian Physiology, pp. 89–111.

35. Davis JK et al. 2010 Evolution of a bitter taste receptor gene cluster in a New World sparrow. Genome Biology and Evolution 2, 358–370. (doi:10.1093/gbe/evq027)

36. Avery ML, Schreiber CL, Decker DG. 1999 Fruit sugar preferences of House Finches. Wilson Ornithological Society 111, 84–88.

37. Lotz CN, Nicolson SW. 1996 Sugar preferences of a nectarivorus passerine bird, the Lesser Double-Collared Sunbird (*Nectarinia Chalybea*). Functional Ecology 10, 360. (doi:10.2307/2390284)

38. Schondube JE, del Rio CM. 2003 Concentration-dependent sugar preferences in nectar-feeding birds: mechanisms and consequences. Functional Ecology 17, 445–453. (doi:10.1046/j.1365-2435.2003.00749.x)

39. Matson KD, Millam JR, Klasing KC. 2001 Thresholds for sweet, salt, and sour taste stimuli in cockatiels (*Nymphicus hollandicus*). Zoo Biology 20, 1–13. (doi:10.1002/zoo.1001)

40. Drewnowski A, Mennella JA, Johnson SL, Bellisle F. 2012 Sweetness and food preference. The Journal of Nutrition, 1142–1148. (doi:10.3945/jn.111.149575.1142S)

41. Bartholomew GA, Cade TJ. 1958 Effects of sodium chloride on the water consumption of house finches. Physiological Zoology 31, 304–310. (doi:10.1086/physzool.31.4.30160936)

42. Duncan CJ. 1960 The sense of taste in birds. Proceedings of the Association of Applied Biologists 4, 409–414.

43. van Dongen MV, van den Berg MC, Vink N, Kok FJ, de Graaf C. 2012 Taste – nutrient relationships in commonly consumed foods. British Journal of Nutrition 108, 140–147. (doi:10.1017/S0007114511005277)

44. Shochat E, Warren PS, Faeth SH, Mcintyre NE, Hope D. 2007 From patterns to emerging processes in mechanistic urban ecology. Trends in Ecology & Evolution 21, 186–191. (doi:10.1016/j.tree.2005.11.019)

45. Auman HJ, Bond AL, Meathrel CE, Richardson AMM. 2011 Urbanization of the Silver Gull: evidence of anthropogenic feeding regimes from stable isotope analyses. Waterbirds 34, 70–76.

46. Coogan SCP, Raubenheimer D, Zantis SP, Machovsky-Capuska GE, Coogan, Sean C. P. Raubenheimer D, Zantis SP, Machovsky-Capuska GE. 2018 Multidimensional nutritional ecology and urban birds. Ecosphere 9, e02177. (doi:10.1002/ecs2.2177)

47. Kirby R, Johnson HE, Alldredge MW, Pauli JN. 2019 The cascading effects of human food on hibernation and cellular aging in free-ranging black bears. Scientific Reports 9, 2197. (doi:10.1038/s41598-019-38937-5)

48. Newsome D, Rodger K. 2008 To feed or not to feed: a contentious issue in wildlife tourism. Australian Zoologist 34, 255–270. (doi:10.7882/fs.2008.029)

49. Schulte-Hostedde AI, Mazal Z, Jardine CM, Gagnon J. 2018 Enhanced access to anthropogenic food waste is related to hyperglycemia in raccoons (*Procyon lotor*). Conservation Physiology 6, 1–6. (doi:10.1093/conphys/coy026)

50. Heiss RS, Clark AB, Mcgowan KJ. 2009 Growth and nutritional state of American Crow nestlings vary between urban and rural habitats. Ecological Applications 19, 829–839. (doi:10.1890/08-0140.1)

51. Kreeger TJ, Waiser MM. 1984 Carpometacarpal deformity in giant Canada Geese (*Branta canadensis maxima* Delacour). Journal of Wildlife Diseases 20, 245–248.

52. Ishigame G, Baxter GS, Lisle AT. 2006 Effects of artificial foods on the blood chemistry of the Australian magpie. In Austral Ecology, pp. 199–207. John Wiley & Sons, Ltd. (doi:10.1111/j.1442-9993.2006.01580.x)

53. Grant PR. 1991 Natural selection and Darwin’s finches. Scientific American 265, 82–87. (doi:10.2307/24938761)

54. Grant PR, Grant RB. 2008 How and Why Species Multiply: The Radiation of Darwin’s Finches. Princeton, New Jersey, USA: Princeton University Press.

55. Hendry AP, Grant PR, Grant RB, Ford HA, Brewer MJ, Podos J. 2006 Possible human impacts on adaptive radiation: beak size bimodality in Darwin’s finches. Proceedings of the Royal Society B: Biological Sciences 273, 1887–1894. (doi:10.1098/rspb.2006.3534)

56. Harvey JA, Chernicky K, Simons SR, Verrett TB, Chaves JA, Knutie SA. 2021 Urban living influences the nesting success of Darwin’s finches in the Galápagos Islands. Ecology and Evolution (doi:10.1002/ece3.7360)

57. De León LF, Bermingham E, Podos J, Hendry AP. 2010 Divergence with gene flow as facilitated by ecological differences: within-island variation in Darwin’s finches. Philosophical Transactions of the Royal Society B-Biological Sciences 365, 1041–1052. (doi:10.1098/rstb.2009.0314)

58. Skelhorn J, Rowe C. 2006 Predator avoidance learning of prey with secreted or stored defences and the evolution of insect defences. Animal Behaviour 72, 827–834. (doi:https://doi.org/10.1016/j.anbehav.2005.12.010)

59. Hämäläinen L, Rowland HM, Mappes J, Thorogood R. 2017 Can video playback provide social information for foraging blue tits ? PeerJ 5, e3062. (doi:10.7717/peerj.3062)

60. Hämäläinen L, Mappes J, Thorogood R, Valkonen JK, Karttunen K, Salmi T, Rowland HM. 2020 Predators’ consumption of unpalatable prey does not vary as a function of bitter taste perception. Behavioral Ecology 31, 383–392. (doi:10.1093/beheco/arz199)

61. Cuthill I, Witter M, Clarke L. 1992 The function of bill-wiping. Animal Behaviour 43, 103–115. (doi:https://doi.org/10.1016/S0003-3472(05)80076-4)

62. Teichmann M, Thorogood R, Hämäläinen L. 2020 Seeing red? Colour biases of foraging birds are context dependent. Animal Cognition 23, 1007–1018. (doi:10.1007/s10071-020-01407-x)

63. Bates DM, Maechler M, Bolker BM, Walker S. 2015 Fitting linear mixed-effects models using lme4. Journal of Statistical Software 67, 1–48.

64. Komsta L, Novomestky F. 2015 MOMENTS: Moments, cumulants, skewness, kurtosis and related tests.

65. Hartig F. 2020 DHARMa: Residual Diagnostics for Hierarchical (Multi-Level / Mixed) Regression Models.

66. Watkins G, Cruz F. 2007 Galapagos at risk: a socioeconomic analysis of the situation in the archipelago.

67. Fitter J, Fitter D, Hosking D. 2016 Wildlife of the Galápagos. 2nd edn. Princeton, New Jersey: Princeton University Press.

68. Roque-Albelo L, Schroeder FC, Conner WE, Bezzerides A, Hoebeke ER, Meinwald J, Eisner T. 2002 Chemical defense and aposematism: the case of Utetheisa galapagensis. Chemoecology 157, 153–157. (doi:10.1007/s00012-002-8341-6)

69. Bravo C, Bautista LM, Garci M, Blanco G, Alonso JC. 2014 Males of a strongly polygynous species consume more poisonous food than females. PLoS One 9, e111057. (doi:10.1371/journal.pone.0111057)

70. Shuker KPN. 2001 The Hidden Power of Animals. Readers Digest.

