## Supplemental Materials for "Darwin’s small and medium ground finches might have taste preferences, but not for human foods"

<u>Assessing latent colour preferences before conducting experimental trials</u>

***Methods:*** To examine whether latent colour preferences could affect our experimental trials, we first presented birds with unflavoured pastry in each of the coloured cups. The method of presentation and data recording was otherwise identical to the experimental trials (see Methods in the manuscript). We then investigated mean differences in feeding events at each location, and for each of the two species that we examined in the experiment. We counted each feeding event from each coloured cup and summed these for each trial.

The number of trials where each species participated varied across locations, with experiments at some locations failing to attract birds in at least 10 trials (Supplementary Table 1). Therefore, we used Cummings estimation plots (using the dabestr package; (Ho et al. 2019)) to first assess evidence for significant differences among the colours at each location (based on 95% confidence intervals, produced using nonparametric bootstrap resampling) separately for each species. 'Pink' was used as the reference level for comparisons as this cup colour contained unflavoured pastry during the taste preference trials. We then used generalised linear mixed effects models to test for differences in colour preferences overall. Here, feeding events were modelled using a negative binomial distribution to account for overdispersion, with cup colour and location included as fixed effects and trial included as a random effect. Data for the two species were then combined to account for small sample sizes at some locations, so species was included as a fixed effect in the model with both species.

***Results:*** For medium ground finches (Supplemental Figure 3), we found a tendency for birds to feed more often from the green and yellow cups, compared to the pink cup, but only at trials conducted at town sites. Although the 95% confidence intervals did not include 0 (green: 0.93 – 11.50; yellow: 2.32 – 12.20), the sampling distributions of the mean differences did. For small ground finches (Supplemental Figure 4), we found a tendency for birds to show a preference towards yellow and blue cups at town sites, and away from green at remote sites, but the number of trials where feeds occurred was very small (N = 5 compared to N = 3 for trials with feeds from pink cups in town, and N = 2 vs. N = 1 at remote sites). When we combined the data from both species, we found no significant differences in feeding preferences across locations (cup colour * location: $\chi^2 = 14.245$, d.f. = 8, p = 0.076) or in the number of feeds from any of the cup colours (cup colour: $\chi^2 = 4.347$, d.f. = 4, p = 0.36) although there was a weakly significant preference to feed from the yellow cup when compared to the pink cup (Supplemental Table 2; versus green: estimate = -0.227 ± 0.217, z = 1.043, p = 0.30) and ground finches fed less often at trials in town sites (Supplemental Table 2). There was no significant difference between species in the number of feeding events (Supplemental Table 2).

Supplemental Table 1. Number of (a) trials (10 minute length) where a given species came to the 3experimental set-up (some trials included both species)

| (a) | Remote | Beach | Town | Total |
|---|---|---|---|---|
| **Taste trials** | | | | |
| Small Ground Finch | 10 | 15 | 14 | 39 |
| Medium Ground Finch | 16 | 4 | 10 | 30 |
| Total | 16 | 15 | 18 | 49 |
| **Colour trials** | | | | |
| Small Ground Finch | 5 | 15 | 9 | 29 |
| Medium Ground Finch | 16 | 5 | 16 | 37 |
| Total | 17 | 17 | 19 | 53 |

Supplemental Table 2. Mean differences (± S.E.) in the number of feeding events by medium and small ground finches (data combined due to small sample sizes, see Supplemental Table 1) from coloured cups containing unflavoured pastry. Differences were estimated using generalised linear mixed effects models with a negative binomial error distribution where trial was included as a random effect. ‘Pink’ is used as the intercept because this cup colour was the reference level in experimental trials with flavoured pastry.

| | Mean difference ± S.E. | Z | p |
|---|---|---|---|
| Intercept (pink, town) | 1.704 ± 0.238 | 7.171 | < 0.001 |
| Blue | 0.262 ± 0.249 | 1.054 | 0.292 |
| Green | 0.234 ± 0.237 | 0.987 | 0.320 |
| Purple | 0.381 ± 0.236 | 1.613 | 0.107 |
| Yellow | 0.461 ± 0.235 | 1.957 | 0.050 |
| Location (beach) | 0.491 ± 0.240 | 2.048 | 0.041 |
| Location (remote) | 0.423 ± 0.219 | 1.921 | 0.054 |
| Species | 0.094 ± 0.200 | 0.471 | 0.638 |

Supplemental Table 3. Mean differences (± S.E.) in the number of feeding events by medium and small ground finches (data combined due to small sample sizes, see Supplemental Table 1) from pastry laced with 'human-food' flavours as compared to neutral unflavoured pastry and a bitter flavoured control. Differences were estimated using generalised linear mixed effects models with a negative binomial error distribution where species nested within trial was included as a random effect and species was included as a fixed effect. The number of feeding events by ground finches on neutral pastry at Town sites is presented here as the intercept.

| | Mean difference ± S.E. | Z | p |
|---|---|---|---|
| Intercept (neutral, town) | 2.178 ± 0.291 | 7.490 | < 0.001 |
| Oily (blue) | 0.136 ± 0.366 | 0.372 | 0.710 |
| Bitter (green) | -0.465 ± 0.360 | -1.292 | 0.196 |
| Sweet (purple) | 0.478 ± 0.416 | 1.148 | 0.251 |
| Salty (yellow) | -0.600 ± 0.364 | -1.650 | 0.099 |
| Location (beach) | 0.664 ± 0.432 | 1.538 | 0.124 |
| Location (remote) | 0.049 ± 0.349 | 0.141 | 0.888 |
| Species | -0.102 ± 0.163 | -0.625 | 0.532 |
| Oily : Beach | -0.357 ± 0.583 | -0.612 | 0.541 |
| Bitter : Beach | 0.502 ± 0.567 | 0.886 | 0.376 |
| Sweet : Beach | -0.689 ± 0.611 | -1.128 | 0.259 |
| Salty : Beach | 0.132 ± 0.558 | 0.237 | 0.813 |
| Oily : Remote | -0.903 ± 0.487 | -1.855 | 0.064 |
| Bitter : Remote | 0.776 ± 0.464 | 1.672 | 0.095 |
| Sweet : Remote | -0.056 ± 0.513 | -0.110 | 0.913 |
| Salty : Remote | 0.545 ± 0.472 | 1.155 | 0.248 |

Supplemental Table 4. Mean differences (± S.E.) in the proportion of feeding events followed by beak-wiping by medium and small ground finches (data combined due to small sample sizes, see Supplemental Table 1) after consuming pastry laced with ‘human-food’ flavours compared to neutral unflavoured pastry and a bitter flavoured control. Differences were estimated using generalised linear mixed effects models with a binomial error distribution where species nested within trial was included as a random effect and species was included as a fixed effect. The proportion of times beak-wiping occurred by medium ground finches on neutral pastry at Town sites is presented here as the intercept.

| | Mean difference ± S.E. | Z | p |
|---|---|---|---|
| Intercept (neutral, town) | -2.362 ± 0.336 | -7.032 | < 0.001 |
| Oily (blue) | 0.551 ± 0.371 | 1.485 | 0.138 |
| Bitter (green) | 0.419 ± 0.431 | 0.972 | 0.331 |
| Sweet (purple) | 0.741 ± 0.379 | 1.955 | 0.051 |
| Salty (yellow) | -0.435 ± 0.517 | -0.842 | 0.400 |
| Location (beach) | 0.690 ± 0.423 | 1.629 | 0.103 |
| Location (remote) | -0.412 ± 0.439 | -0.939 | 0.348 |
| Species | 0.528 ± 0.198 | 2.672 | 0.008 |
| Oily : Beach | -1.307 ± 0.559 | -2.336 | 0.020 |
| Bitter : Beach | -1.121 ± 0.571 | -1.964 | 0.050 |
| Sweet : Beach | -1.413 ± 0.545 | -2.592 | 0.010 |
| Salty : Beach | 0.449 ± 0.632 | 0.711 | 0.477 |
| Oily : Remote | 0.316 ± 0.572 | 0.553 | 0.580 |
| Bitter : Remote | -0.298 ± 0.574 | -0.519 | 0.604 |
| Sweet : Remote | 0.563 ± 0.509 | 1.108 | 0.268 |
| Salty : Remote | 1.346 ± 0.633 | 2.128 | 0.033 |

Supplemental Figure 1. Locations and representative photographs of the three different sites experimental trials were conducted. (A) is the remote site with no human foods, (B) is the beach site which has human foods from picnics, and (C) is the town site where human foods are abundant. (D) shows a map of the sites where experiments were conducted and match the letters in this legend.

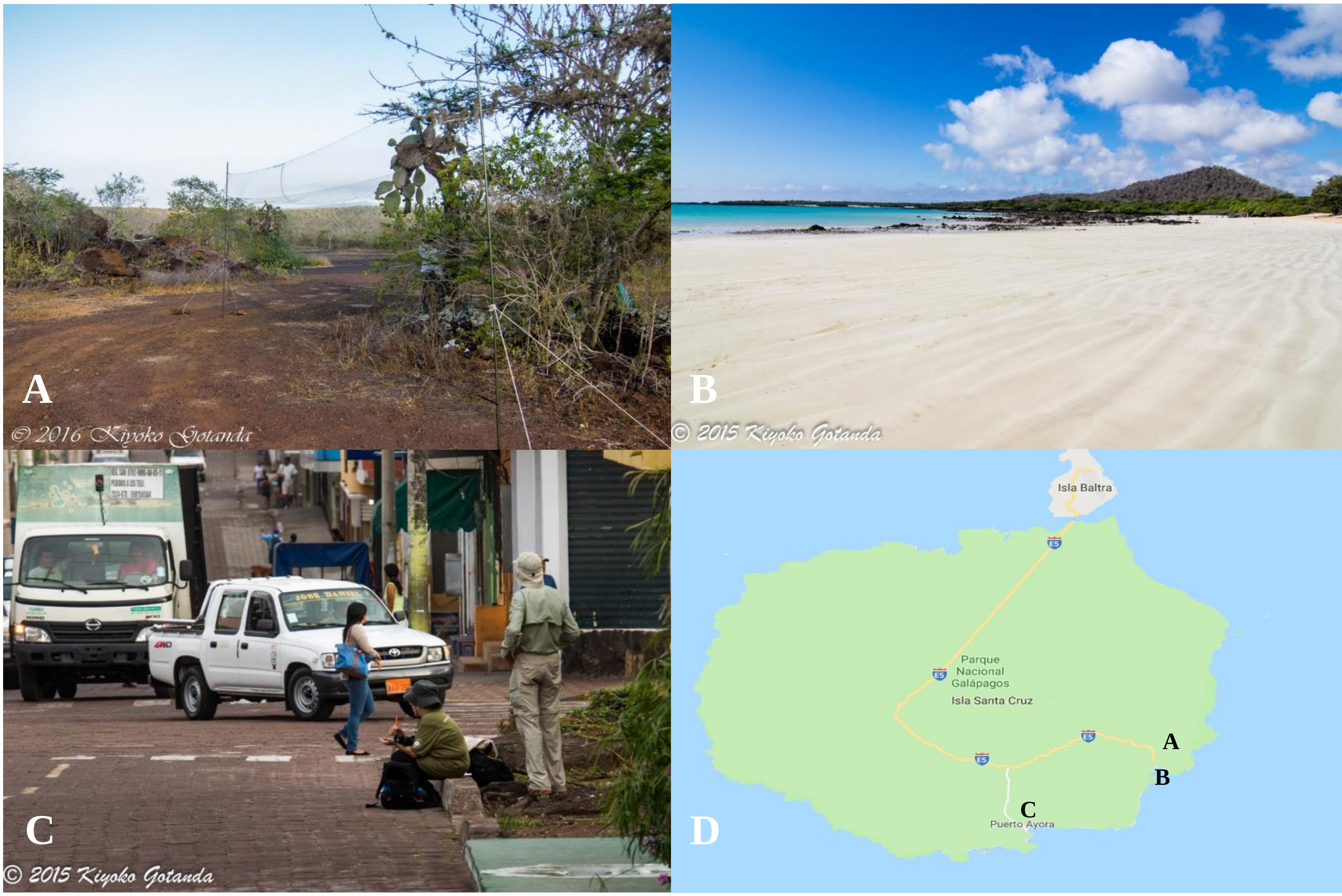

Supplemental Figure 2. Photo showing a finch feeding from a cup during one of the trials.

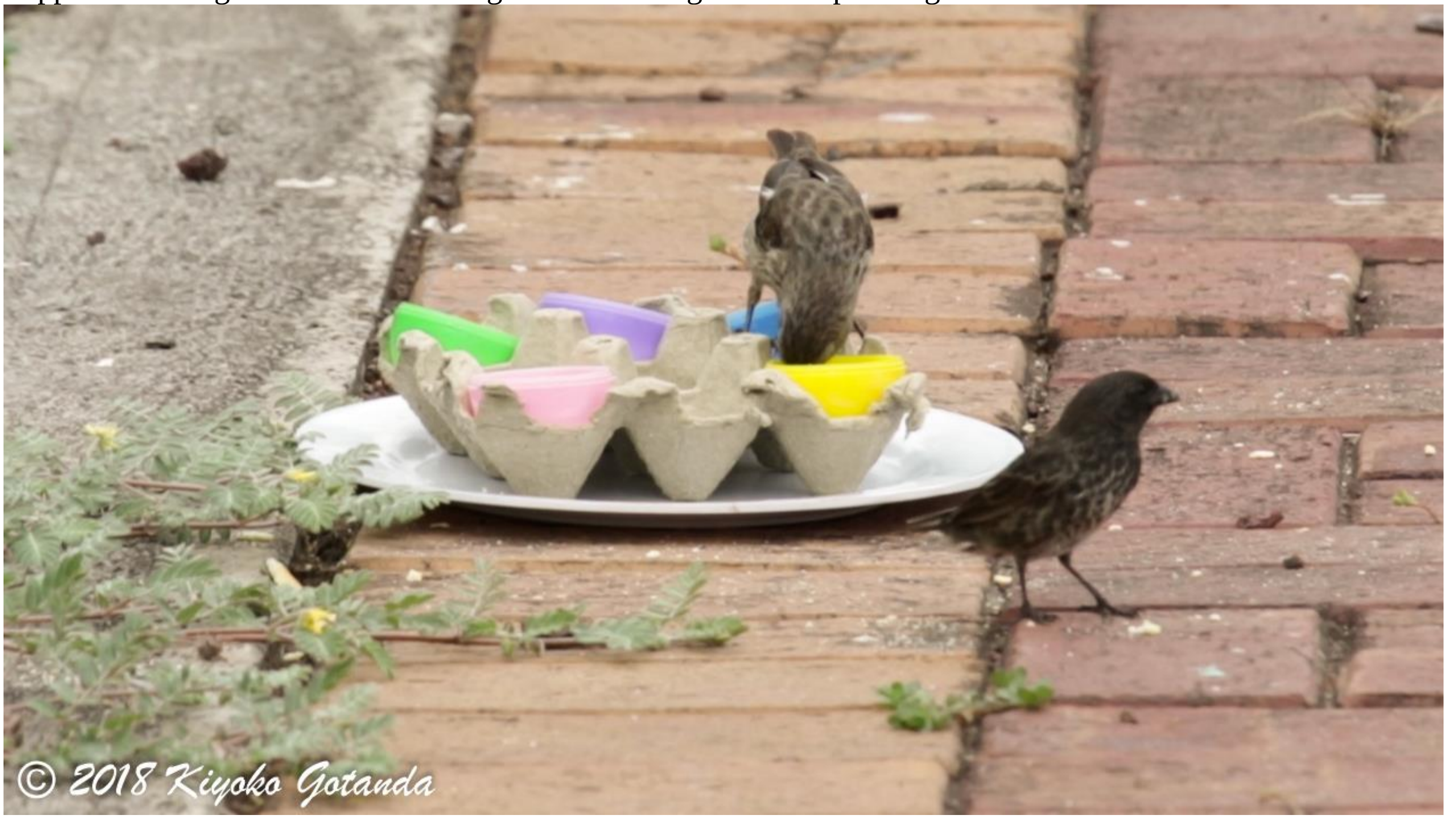

Supplemental Figure 3: The mean difference in feeding events by medium ground finches when presented with unflavoured pastry in coloured cups at sites in remote or town locations (there were no feeding events to the comparison ‘pink’ cup at beach locations). The raw data are plotted in **(top)** with the number of trials indicated below the x-axis. The mean difference between each coloured cup and the ‘pink’ cup (black filled circle) is plotted in **(bottom)** with its bootstrap sampling distribution (grey shaded area). The 95% confidence interval for each difference is indicated by black vertical error bars, and the horizontal bar at 0 indicates no mean difference.

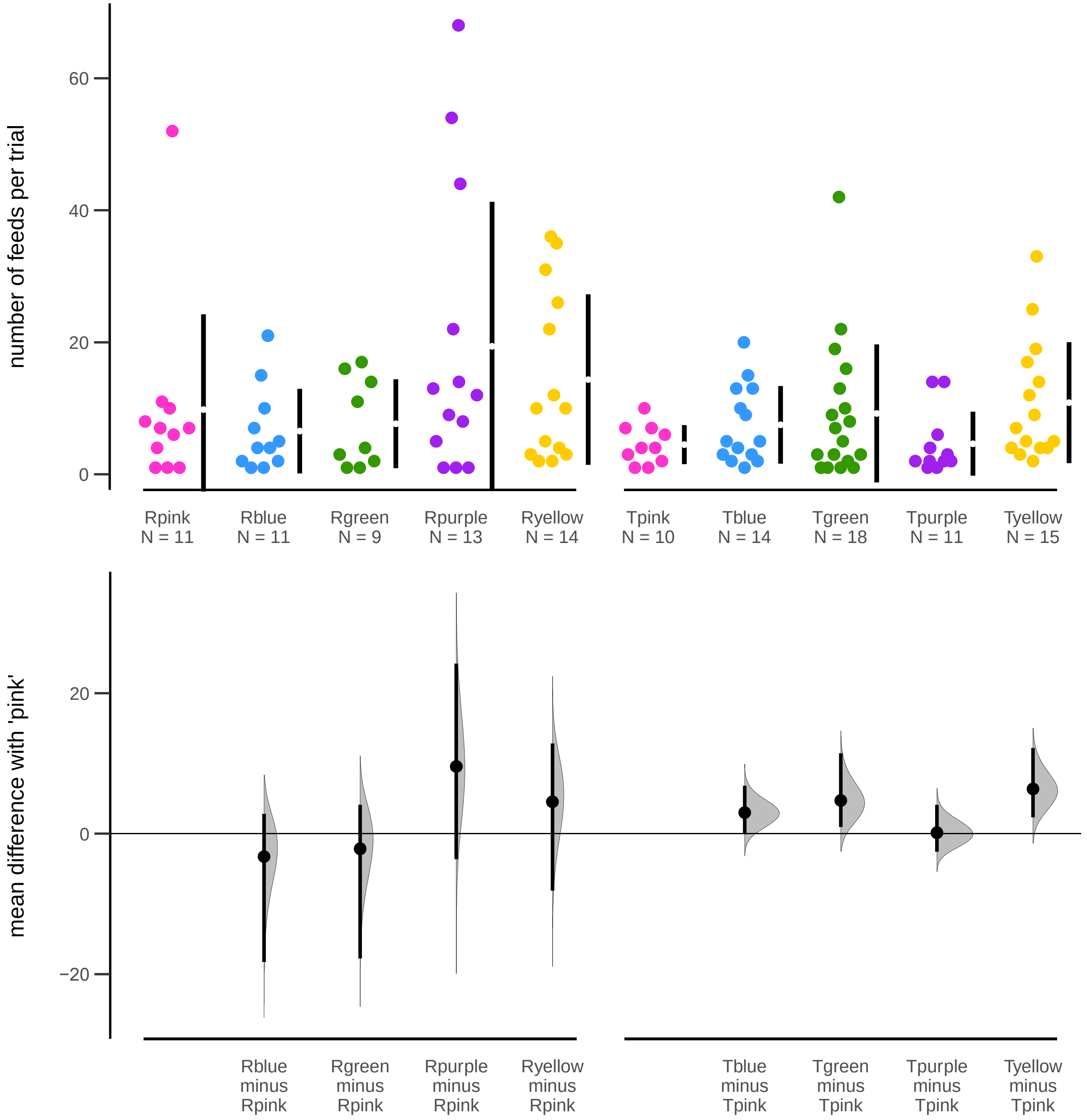

Supplemental Figure 4: The mean difference in feeding events by small ground finches when presented with unflavoured pastry in coloured cups at sites in remote, beach, or town locations. The raw data are plotted in **(top)** with the number of trials indicated below the x-axis. The mean difference between each coloured cup and the 'pink' cup (black filled circle) is plotted in **(bottom)** with its bootstrap sampling distribution (grey shaded area). The 95% confidence interval for each difference is indicated by black vertical error bars, and the horizontal bar at 0 indicates no mean difference.

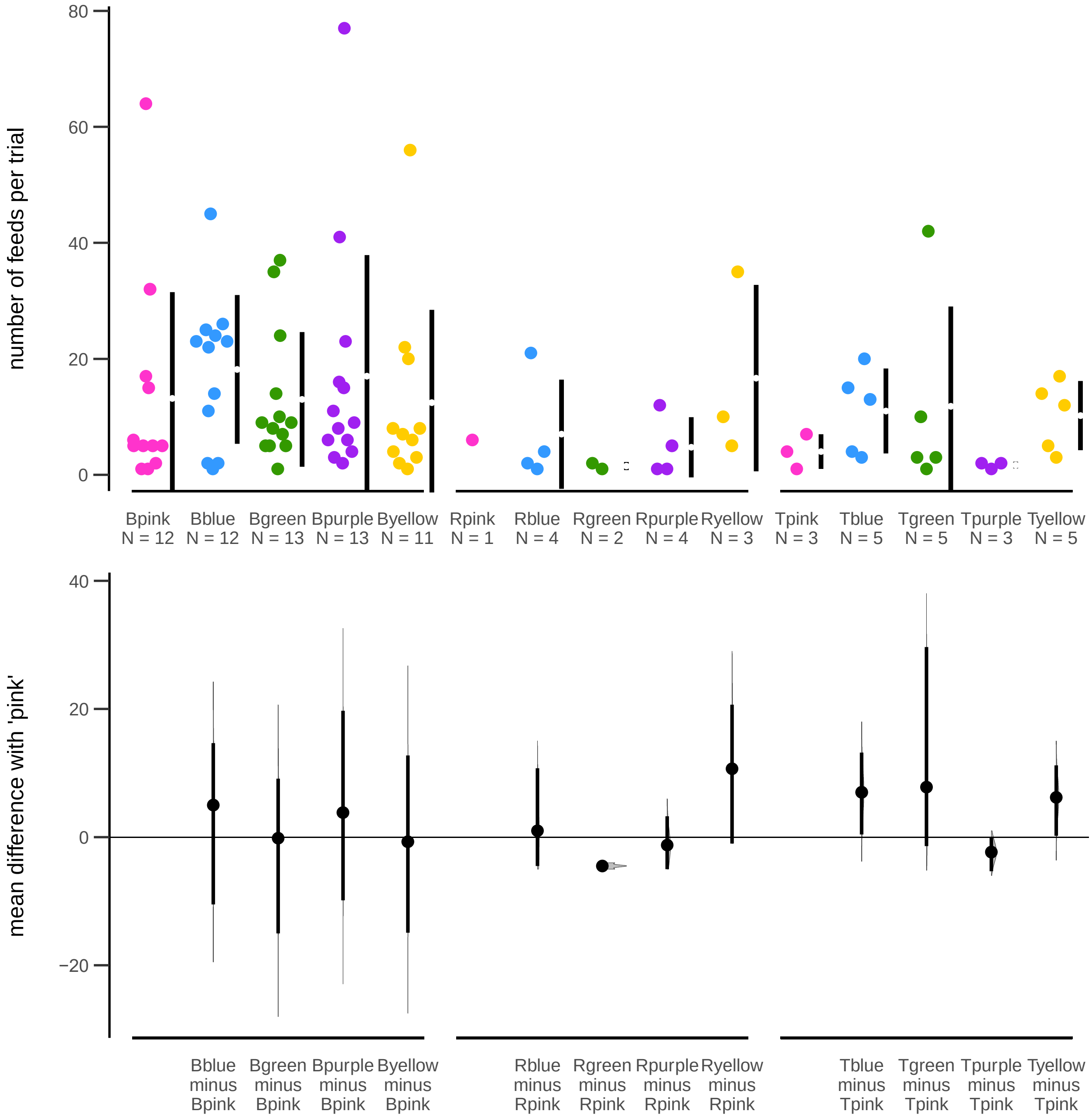

Supplemental Figure 5: Packaging of popular food items available for purchase on the Galapagos Islands.

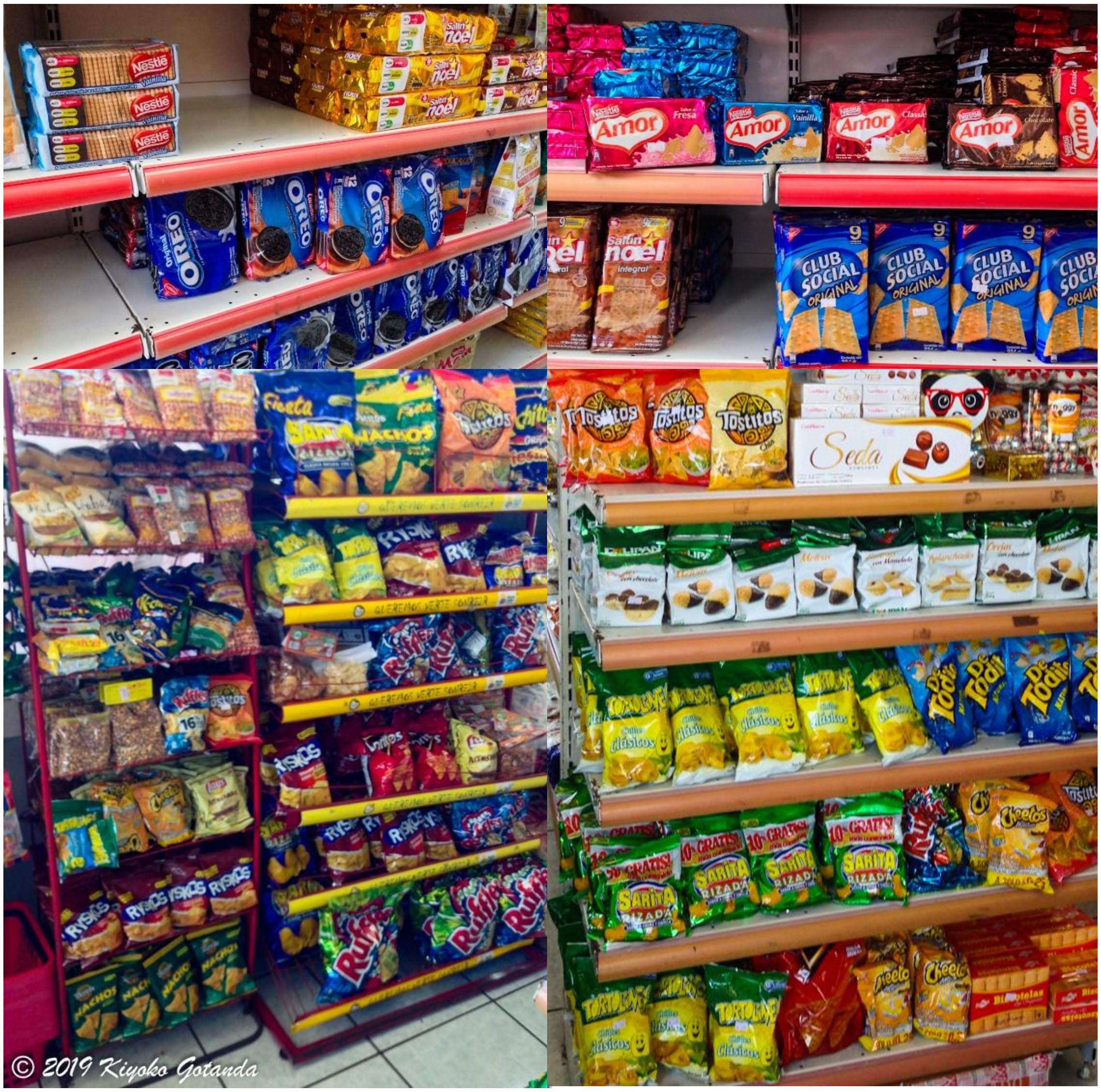
